# APEs: Agentic Prediction of Epitopes to Guide Antibody–Antigen Structure Prediction

**DOI:** 10.64898/2026.11.23.768576

**Authors:** Minji Kang, Min Su Yoon, Jeonghoon Park, Byunghyun Moon, Minkyung Baek

**Affiliations:** Department of Biological Sciences, Seoul National University, Seoul, Republic of Korea; College of Pharmacy, Seoul National University, Seoul, Republic of Korea; Artificial Intelligence Institute, Seoul National University, Seoul, Republic of Korea

**Author notes:** These authors contributed equally; orders of the names are randomized (Minji Kang, Min Su Yoon, Jeonghoon Park).

## Abstract

Antibody-antigen complex prediction remains difficult because the surface recognized by a specific antibody is often uncertain, and model confidence may not distinguish alternative poses. We developed APEs (Agentic Prediction of Epitopes for Structure Prediction), an agentic system that retrieves public evidence, interprets predicted antibody and antigen structures, and produces ranked residue-level epitope hypotheses. For retrospective evaluation, web retrieval was restricted to sources released before each CASP target entry date. Across eight CASP16 and six released CASP17 targets, the top-ranked patch reached residue precision of at least 0.5 for nine of 14 targets. We then used the predicted epitopes to steer additional Protenix-v1 sampling and to compare steered finalists with the highest ipSAE unsteered model. The retrospective best selection across three independent router calls increased mean DockQ from 0.306 to 0.375; DockQ increased from 0.089 to 0.679 for H2321 and from 0.005 to 0.379 for H2346. These results show that publicly available evidence specific to the target antibody–antigen complex can serve as an inference-time signal for epitope localization, guided complex sampling, and model selection. We also release predictions for CASP17 targets whose experimental structures had not been released when the predictions were finalized, providing a prospective test of APEs (https://github.com/CSSB-SNU/ape).

## Introduction

Recent advances in protein structure prediction have substantially improved protein–protein complex modeling^1–3^. However, antibody–antigen complexes remain particularly challenging, with CASP assessments and independent benchmarks reporting lower success rates than for other protein–protein complexes^4–6^.

One challenge is that a particular antibody and its antigen generally lack the paired evolutionary information that can help identify interchain contacts. The sequence diversity and conformational variability of antibody CDRs further complicate prediction. Near-native models can still appear within sampled ensembles, but confidence scores do not always rank them first^5,7^. Information about the antigen surface recognized by an antibody can therefore provide a useful constraint, even when it does not determine the complete binding orientation.

Epitope and contact restraints can improve antibody-antigen structure prediction when informative sites are known. Scanning multiple antigen sites can also broaden the sampled binding modes, although antigen-only B-cell epitope rankings did not outperform a random-pocket control^8,9^. However, identifying regions that are generally likely to serve as epitopes is different from determining which surface is recognized by a particular antibody. Antibody-conditioned predictors such as PEPNet^10^ have sought to address this distinction by inferring the epitope from the antibody and antigen sequences and structures provided as input. Nevertheless, antibody-specific epitope localization remains challenging, and EpiBench similarly found that general-purpose LLMs without web access had limited ability to identify the bound epitope from antibody and antigen sequences alone^11^.

Relevant evidence for a given antibody-antigen pair may already exist in mutational studies, hydrogen-deuterium exchange experiments, competition assays, domain-mapping studies, patents, or solved structures of homologous complexes. These observations are distributed across various sources and often use different numbering conventions. Applying them to a new target requires identifying the molecules involved, mapping reported residues to the query sequence, judging the relevance of evidence from homologs or related antibodies, and assessing their relevance to the supplied structures. LLM-based agents that combine literature retrieval with structural analysis offer a way to organize these various pieces of evidence into antibody-specific epitope hypotheses.

Here, we present APEs, an agentic system that brings these steps together. APEs identifies the antigen and antibody, retrieves target-specific evidence, maps reported residues onto the query sequences, and proposes antibody-specific patches on the antigen surface. Three launcher agents generate independent hypotheses, and a Geometric Organizer, a Refiner, and a Judge combine overlapping proposals and rank the retained patches. Each hypothesis includes the proposed residues, predicted antibody-chain contacts, supporting and conflicting evidence, and a written rationale.

We evaluated the predicted patches against experimental interfaces from CASP16 and released CASP17 targets, restricting retrieval by each target’s CASP entry date. We then used the patches to guide additional sampling and asked an evidence-aware router to choose among two steered finalists and an unsteered Protenix-v1 baseline^3^. We also release predictions generated for CASP17 targets whose experimental structures had not been released when the predictions were finalized. The prediction files are available at https://github.com/CSSB-SNU/ape, enabling prospective evaluation as the corresponding structures become public.

## Results

### APEs generates ranked epitope hypotheses from target-specific evidence

APEs first identifies the antibody–antigen pair and links reported evidence to the query sequences (**Fig. 1**). The launcher agents then search independently and propose candidate patches on the antigen surface. The gatherer combines overlapping proposals, reviews the evidence behind them, and ranks the retained hypotheses. The ranked patches are then used to guide structure sampling and to compare the resulting models.

For each retained hypothesis, APEs reports the proposed epitope residues, residue-level contact scores, predicted heavy-chain, light-chain, or combined contact assignments, supporting and conflicting evidence, and a written rationale. Relative patch confidence scores rank alternative epitope regions for the same target, whereas residue-level contact scores prioritize individual residues within each patch. The five highest-scoring residues define the Top5 steering condition used in the subsequent epitope-steered sampling stage.

**Figure 1.**
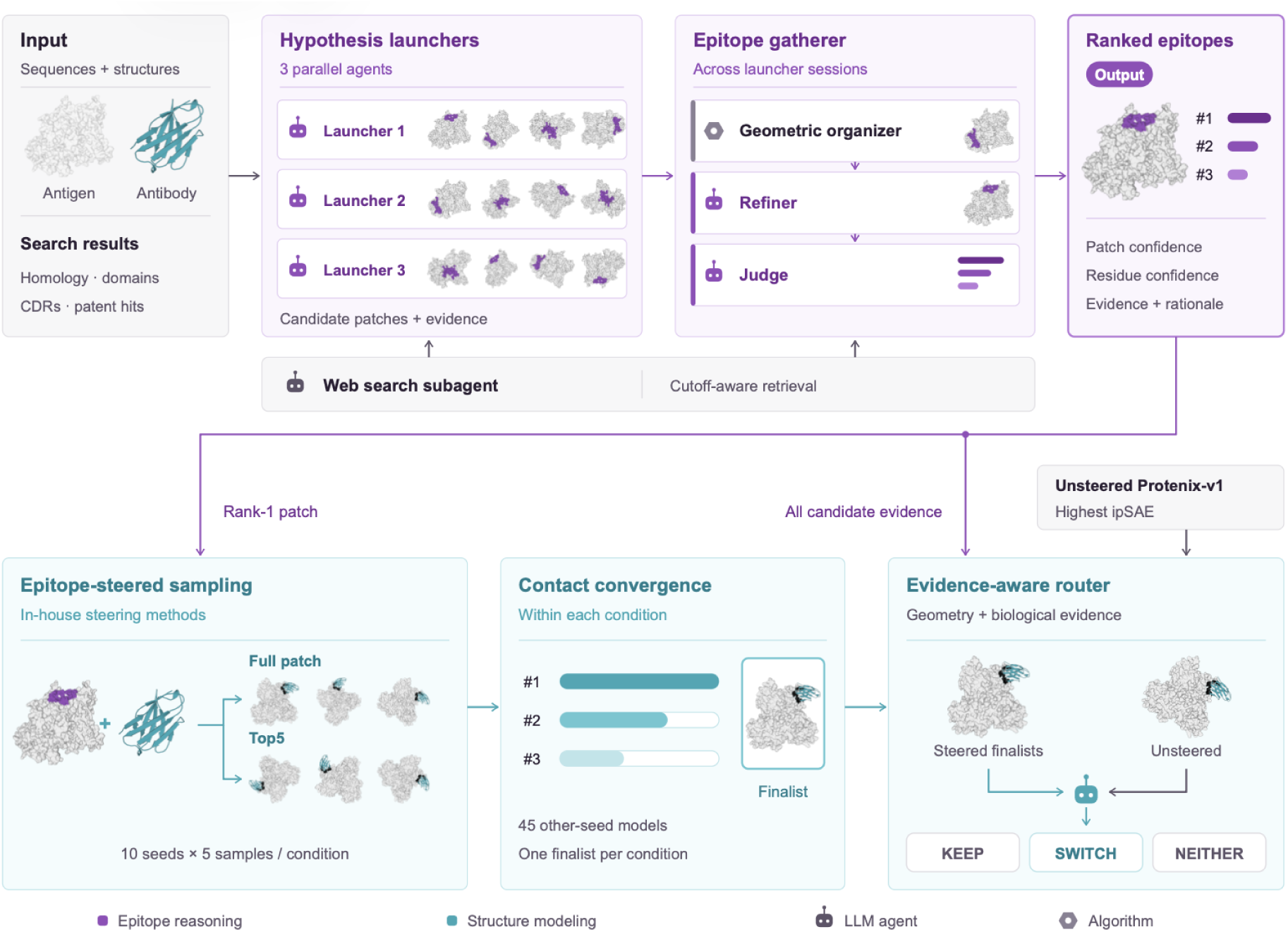
Overview of the APEs pipeline. Parallel agents identify the antibody and antigen, retrieve evidence specific to the target complex, and propose antigen-surface patches as possible epitopes. A geometric organizer, refiner, and judge merge overlapping proposals and rank the predicted patches. The rank-1 branch shown in the diagram is used when relative patch confidence is at least 0.4; otherwise, sampling is repeated for each predicted candidate. An evidence-aware router compares the steered finalists with the highest ipSAE unsteered Protenix-v1 model to choose final models.

### APEs localizes antibody-specific epitope regions in CASP targets

We evaluated APEs on antibody–antigen complex targets from CASP16 and CASP17 for which experimental structures had already been released. The top-ranked patch reached residue precision of at least 0.5 for nine of the 14 targets. When all predicted candidates were considered, APEs proposed at least one patch at or above this threshold for six of eight CASP16 targets and four of six CASP17 targets, giving ten of 14 targets overall (**Fig. 2a**).

Patch confidence tended to track rank-1 accuracy in this cohort. Rank-1 patches with precision of at least 0.5 had a mean patch-confidence score of 0.544, compared with 0.254 for patches below this threshold. The mean score difference between ranks 1 and 2 was 0.254 when the rank-1 patch reached precision of at least 0.5 and 0.082 otherwise (**Fig. 2b**). Residue-level contact scores provided a second ranking within each patch. Among patches with relative patch confidence of at least 0.4, increasing the residue-score threshold generally increased precision, indicating that true contacts tended to receive higher scores than false positives. Low-confidence patches remained imprecise across residue-score thresholds (**Fig. S1**).

**Figure 2.**
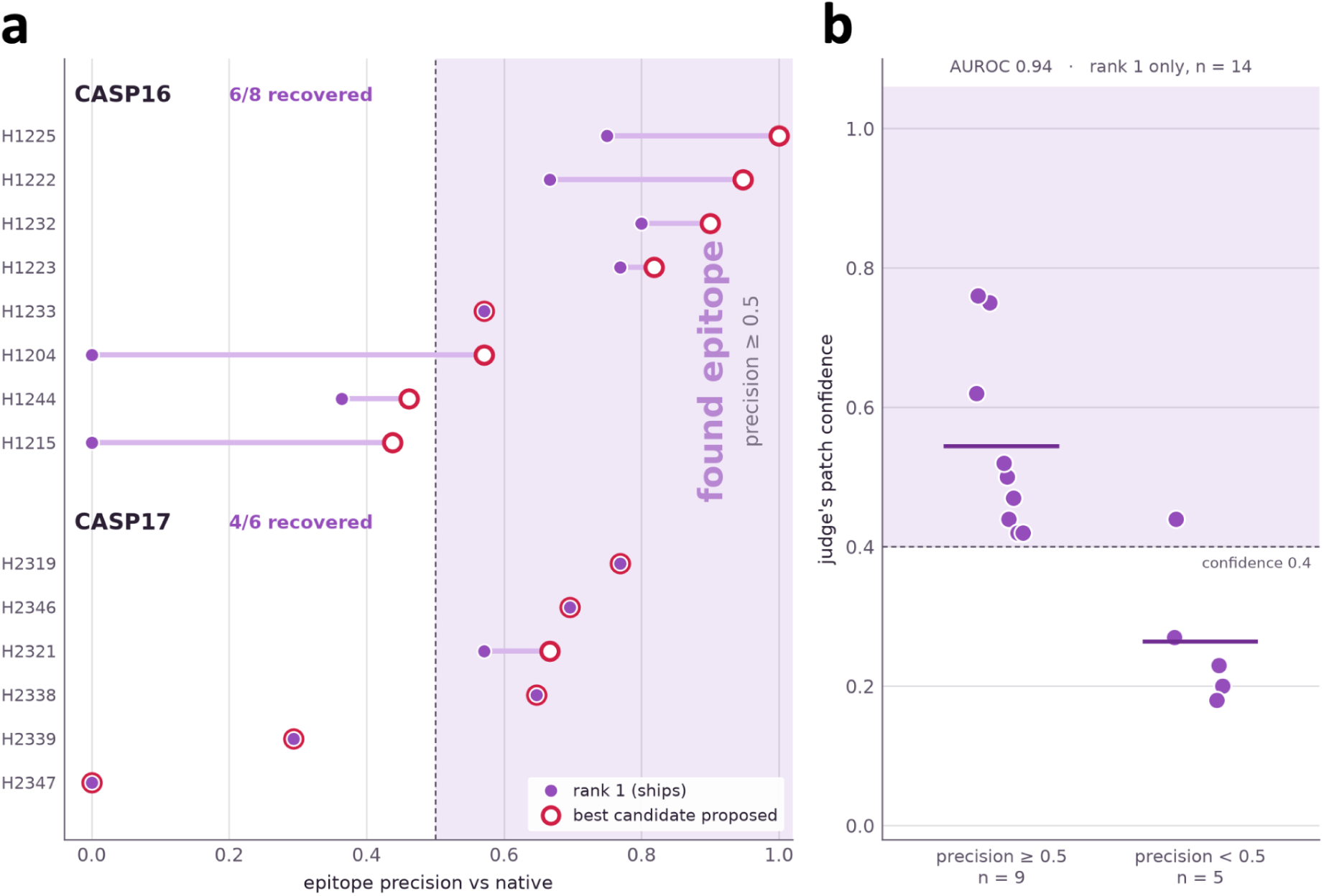
Epitope localization and relative patch confidence on released CASP16 and CASP17 targets. **a**, Residue-level precision of the rank-1 patch and the highest-precision candidate among all predicted patches for each target. The reference epitope was derived from the experimental antibody–antigen complex: an antigen residue was defined as an epitope residue if any of its heavy atoms was within 5 Å of any heavy atom in the antibody. **b,** Relative patch-confidence scores for rank-1 patches grouped by whether their precision was ≥ 0.5 or < 0.5.

### APEs maps evidence at different resolutions to antigen surfaces

**Figure 3.**
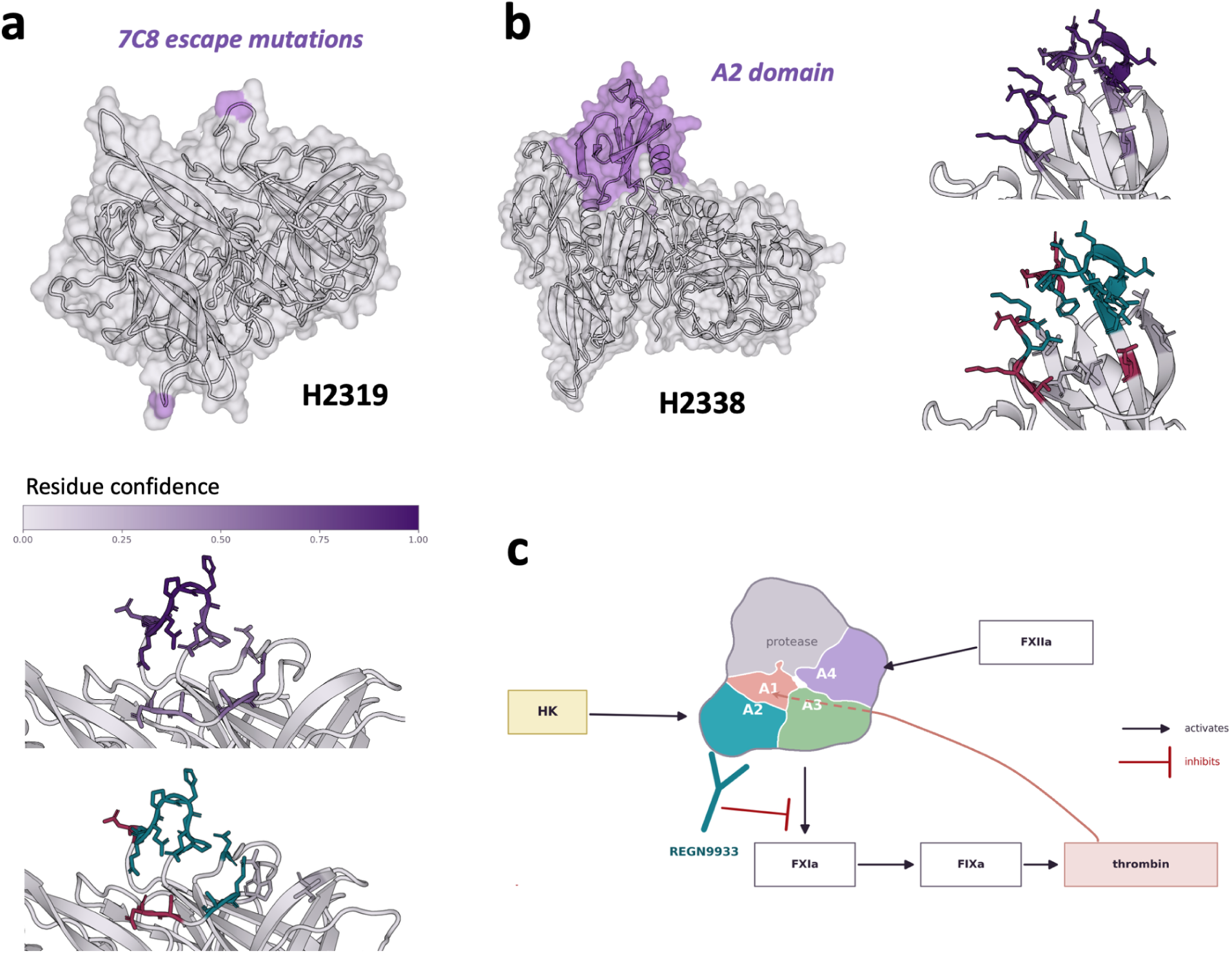
Evidence-guided epitope localization for H2319 and H2338. **a**, For H2319, the reported 7C8 escape substitutions Q542R and P543L were mapped to Q136 and P137 in the query sequence, anchoring the predicted epitope on two symmetry-related sites of the human astrovirus VA1 capsid spike. The close-up views show the top-ranked patch colored by residue-level contact score (middle) and its overlap with the reference epitope (bottom). **b,** For H2338, antibody-specific evidence localized REGN9933 binding to the A2 apple domain of factor XI, shown in purple. APEs combined this domain-level constraint with evidence from other A2-directed antibodies and structural analysis to select a surface within the domain. The close-up views show the residue-level contact scores and the overlap between the predicted and reference epitopes. In the overlap views in **a** and **b**, true-positive residues are shown in teal, false-positive residues in red, and reference epitope residues missed by APEs in gray. **c,** Proposed functional consequence of REGN9933 binding at the A2 domain. High-molecular-weight kininogen (HK) binds A2 and supports FXIIa-mediated activation of FXI. Occupancy of this region by REGN9933 could inhibit this contact-pathway activation while leaving thrombin-mediated feedback activation and the activity of preformed FXIa intact. HK, high-molecular-weight kininogen; FXIIa, activated factor XII; FXIa, activated factor XI; FIXa, activated factor IX.

The H2319 and H2338 cases illustrate how APEs used evidence at different resolutions. For H2319, antibody-specific escape mutations localized the interface at the residue level. For H2338, the available antibody-specific evidence identified the antigen domain, but evidence from related antibodies and structural compatibility were still needed to select a surface within that domain.

#### Case H2319

APEs identified the antigen as the human astrovirus VA1 capsid spike and the antibody as 7C8. One hypothesis launcher retrieved the reported 7C8 escape substitutions Q542R and P543L^12,13^, mapped them to Q136 and P137 in the query antigen, and expanded a candidate patch around these anchors (**Fig. 3a**). Another hypothesis launcher proposed a similar region from geometric and chemical complementarity without using the escape mutation evidence.

The geometric organizer retained two nearby sets of proposed epitope residues as separate candidates because their combined spatial extent was larger than the antigen surface that the antibody CDRs could plausibly contact in a single binding pose. After refinement, the Judge ranked these candidates first and second, with relative patch-confidence scores of 0.50 and 0.41. Although the candidates differed in their exact residue boundaries, both localized the epitope to the same antigen neighborhood. The top-ranked candidate reached a precision of 0.769.

The predicted patch contained 13 residues on each of the two symmetric antigen chains and localized the correct epitope region, although its boundary was shifted relative to the experimental interface. The residue-level contact scores nevertheless prioritized the true interface contacts within the patch: nine of the ten highest-scoring residues were true contacts, whereas two of the three lowest-scoring residues were false positives. Thus, the residue-wise ranking identified a more accurate contact subset within a correctly localized but imperfectly delineated patch.

#### Case H2338

APEs identified H2338 as a complex between human coagulation factor XI (FXI) and the REGN9933 Fab^14^. All three launchers matched the antibody heavy-chain sequence to a patent that localized REGN9933 binding to the 84-residue A2 apple domain of the 607-residue antigen but did not resolve the epitope at the residue-level resolution^15^. Other A2-directed antibodies protected residues 133–148 and 151–163 by HDX-MS^16^. APEs combined this indirect evidence with solvent exposure and chemical complementarity to rank a surface within the A2 domain (**Fig. 3b**). The top-ranked patch received a relative confidence score of 0.52 and reached precision and recall of 0.647.

APEs further used the predicted epitope to propose a functional mechanism for REGN9933 (**Fig. 3c**). FXI can be activated through two routes. In the contact pathway, high-molecular-weight kininogen (HK) binds the A2 apple domain and presents FXI for cleavage by FXIIa. In a separate feedback pathway, thrombin activates FXI directly and helps sustain normal haemostasis. The predicted REGN9933-binding surface was positioned at the HK-binding DFP-motif pocket on A2, suggesting that Fab occupancy could disrupt FXI–HK association and selectively reduce FXIIa-mediated activation. Because this mechanism does not target thrombin-mediated activation or the catalytic activity of already formed FXIa, it would inhibit contact-pathway activation without completely disabling FXI function. Indeed, APEs also referenced the previously reported mechanisms of the A2-directed antibodies 14E11 and AB023 to support this hypothesis^17–19^. A subsequently released cryo-EM structure of REGN9933 bound to FXIa (PDB 9YRQ) and associated functional assays support the APEs-predicted binding site and proposed mechanism^20^.

### Epitope hypotheses guide complex sampling and model selection

To test whether APEs’ epitope hypotheses were applicable for complex prediction, we used them both to steer additional structure sampling and to support model selection (**Fig. 4**). For each target, sampling was guided either by the complete predicted patch or by its five highest-scoring residues (Top5). The model with the highest contact convergence from each steering condition was then compared with the highest ipSAE^21^ unsteered baseline by an evidence-aware router, which considered agreement with the epitope evidence, interface geometry, and biological context. Across the 14 targets, mean DockQ^22^ was 0.306 for the unsteered baseline and 0.375 for the retrospective highest DockQ selection among three independent router calls (**Fig. 4a–c**). The cases below illustrate how the epitope evidence and structural context affected individual selections.

**Figure 4.**
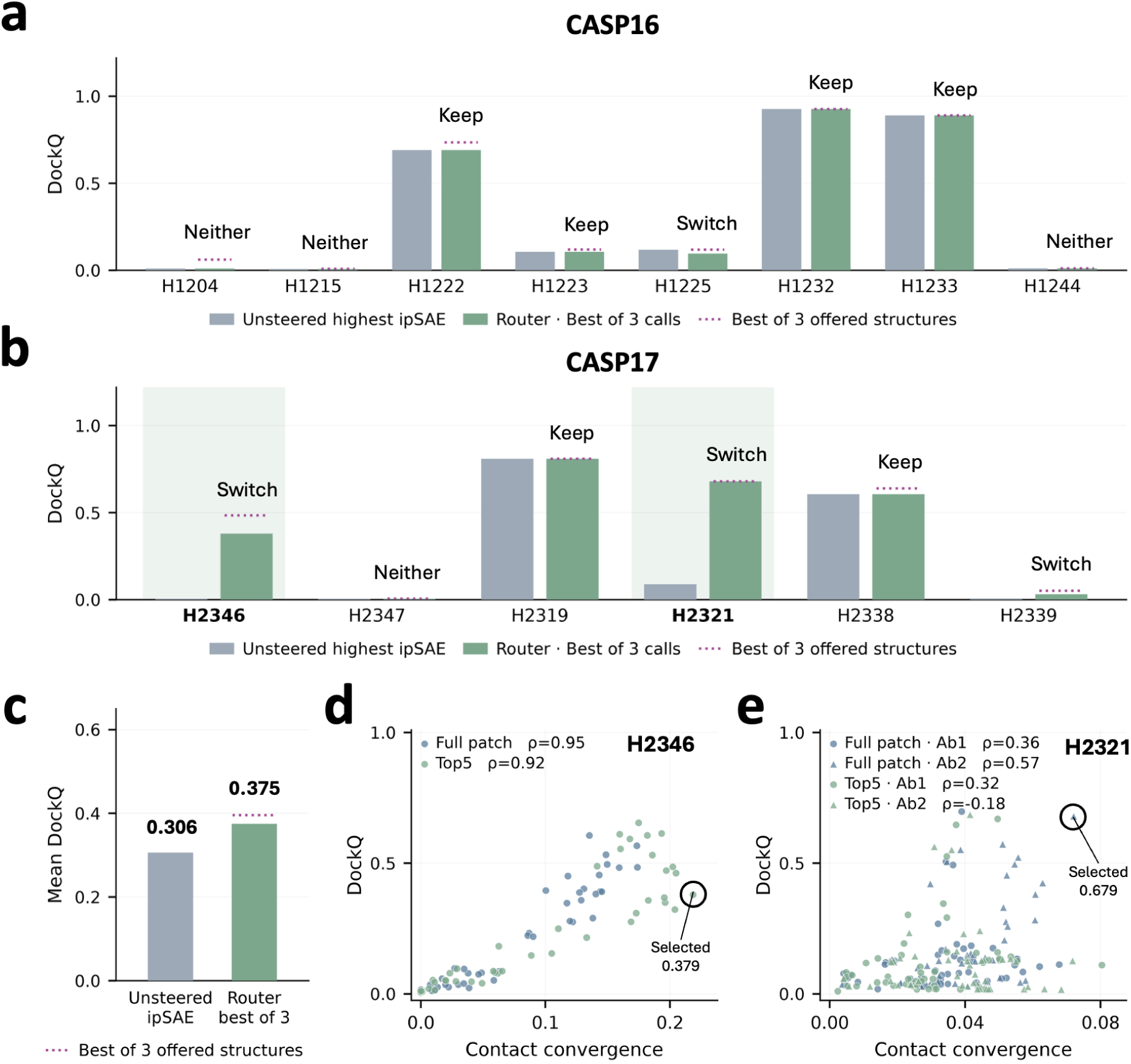
Structure selection using predicted epitopes. **a-b**, Whole-assembly DockQ for the highest ipSAE unsteered baseline and the highest DockQ selection among three independent router calls per target. Dotted lines show the highest DockQ among the three structures offered to the router. **c,** Across 14 targets, mean DockQ was 0.306 for the unsteered baseline and 0.375 for the best router selection. **d-e,** Contact convergence versus DockQ for H2346 and H2321. Each point is one steered structure; rings mark the selected models.

#### Case H2346

H2346 was a Fab–Fab recognition problem between a chimeric Fab and the BCMA-targeted Fab arm of linvoseltamab^23^. Subunits 1 and 2 form the chimeric Fab, whereas subunits 3 and 4 form the linvoseltamab arm. The router was given the retrieved evidence that an anti-κ Fab was present in the linvoseltamab–BCMA cryo-EM sample, supporting an interface at the κ light-chain constant domain of subunit 4^24^. The highest ipSAE unsteered baseline assembled the chains incorrectly, pairing subunit 1 with subunit 4 and subunit 3 with subunit 2 instead of forming the expected subunit 1–2 and subunit 3–4 Fab pairs. A router call explicitly identified this chain swap as a pairing error. Both steered models restored the correct Fab pairs and placed the interface on the evidence-supported κ constant domain. The router distinguished between them based on interface geometry: the full-patch model contained 17 interface heavy-atom clashes, whereas the Top5 model had none. All three router calls therefore selected the Top5 model, which reached a DockQ of 0.379 (**Fig. 4d**, **Fig. 5a**) compared with 0.005 for the baseline. Retrospective evaluation showed that the unselected full-patch model had a higher DockQ of 0.484, so the clash-free selection did not maximize native structural accuracy in this case.

#### Case H2321

H2321 was a complex between the human astrovirus VA1 capsid spike and antibody 2A2^12,13^. Escape mapping implicated two antigen residues in antibody binding. The unsteered baseline had a high ipSAE of 0.710 but contacted only one of these residues (**Fig. 5b**). Among the structures generated using the complete predicted epitope patch as a steering restraint, the model with the highest contact convergence contacted both residues and reached a DockQ of 0.679 (**Fig. 4e**, **Fig. 5b**), compared with 0.089 for the baseline. Two of the three router calls selected this steered model. The remaining call did not treat contact with both escape-mapped residues as decisive because escape mutations can affect binding without being located at the direct interface. Given these uncertainties, the call retained the high-confidence baseline, which had clean interface geometry. The three router calls therefore differed in how they balanced antibody-specific experimental evidence against predictor confidence and interface geometry.

**Figure 5.**
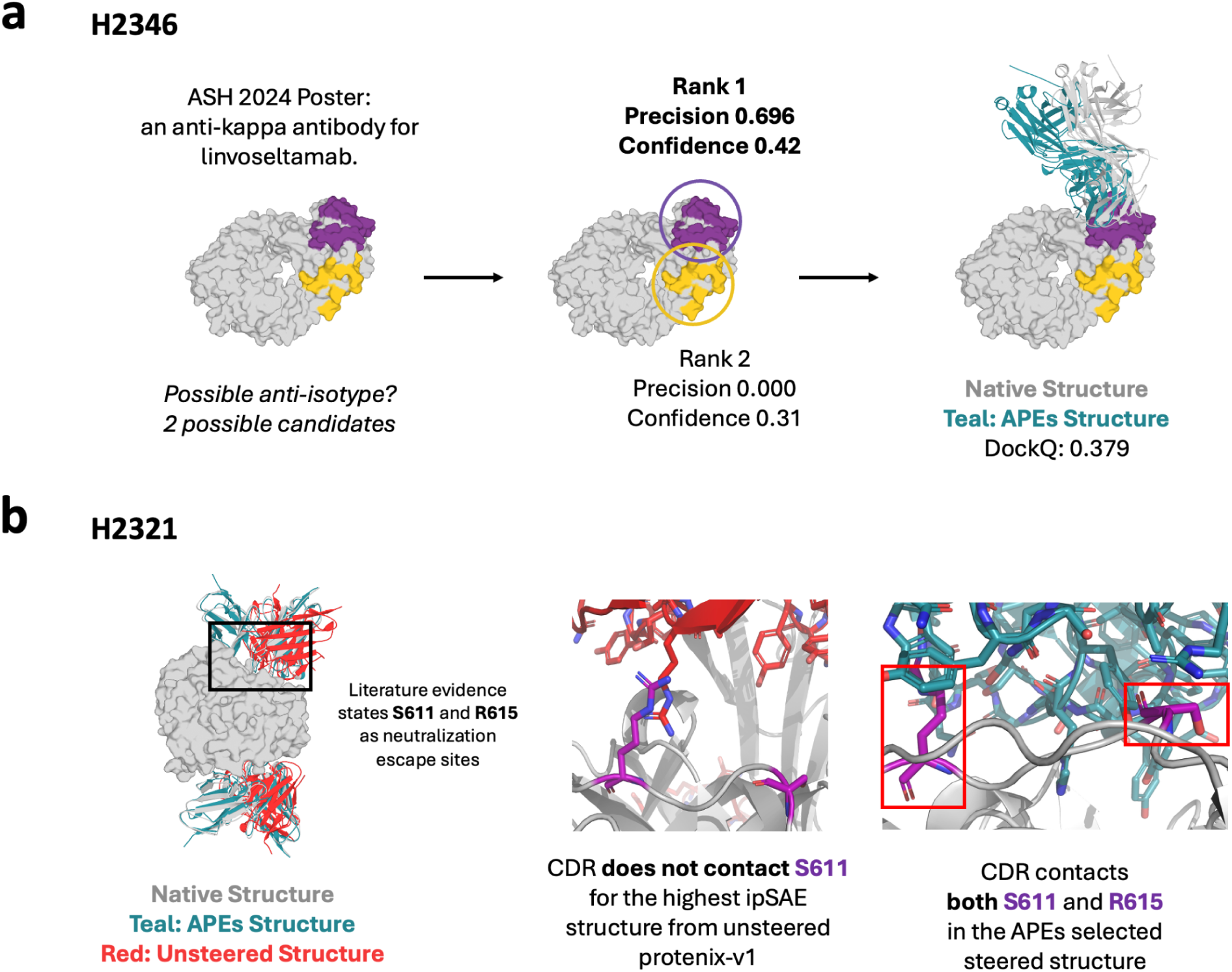
Antibody-specific evidence guides complex modeling for H2346 and H2321. **a**, For H2346, APEs found an ASH 2024 poster describing an anti-κ Fab that bound the same BCMA-targeted Fab arm of linvoseltamab. This evidence shifted the predicted epitope toward the κ light-chain constant domain. The rank-1 patch (purple; relative patch confidence, 0.42) achieved a residue precision of 0.696, whereas the rank-2 patch (yellow; relative patch confidence, 0.31) had no overlap with the native epitope. The router selected a clash-free Top5-steered model (teal) that preserved the correct heavy–light chain pairings and achieved a DockQ of 0.379 against the native assembly (gray). The highest-ipSAE unsteered model is not shown because it contained incorrect heavy–light chain pairings. **b,** H2321 is a complex between the human astrovirus VA1 capsid spike and antibody 2A2. APEs identified S611 and R615 as reported neutralization-escape sites. The highest ipSAE unsteered model (red; ipSAE, 0.710) placed its CDRs in contact with R615 but not S611. The APEs-selected steered model (teal) contacted both escape sites and more closely reproduced the native binding mode (gray), increasing DockQ from 0.089 to 0.679.

#### Case H1225

H1225 was a CASP16 target comprising a 49-residue fragment of the central conserved domain (CCD) of respiratory syncytial virus (RSV) glycoprotein G in complex with the monoclonal antibody Fab 1G1. The rank-1 patch had a precision above 0.5 against the experimental epitope (Fig. 2a), indicating that APEs localized the epitope correctly. This result did not translate into an accurate complex structure after steering. The retrospective best selection across the three router calls had a DockQ of 0.097, while the unsteered baseline had a DockQ of 0.118; both predictions remained inaccurate (**Fig. 4a**). Despite having selected a structure that satisfies all of APEs’ predicted residues, this case exemplifies how accurate knowledge of the epitope does not always translate to a better orientation.

## Discussion

Taken together, these results provide a proof of concept that an agentic system can make dispersed antibody-specific evidence actionable for structure prediction. APEs converts information from publications, patents, sequence searches, and structural analysis into ranked, residue-mapped epitope hypotheses. On the released CASP targets, the rank-1 hypothesis reached the precision threshold for nine of 14 targets, and the candidate pool contained a qualifying patch for ten. Each hypothesis also retained the evidence and rationale behind the proposed region, allowing the prediction to be examined at the level of molecular identity, residue mapping, and competing interpretations.

APEs handled evidence with different levels of specificity. Clone-specific escape mutations provided residue-level anchors for H2319, whereas the evidence for H2338 identified a domain and required information from related antibodies and antigen-surface analysis to select a candidate region. Retaining several hypotheses was useful in the latter setting because the available evidence did not uniquely determine a single surface. Residue-level contact scores added a second layer of resolution within each patch: higher-scoring residues were enriched for true contacts when the patch itself was correctly localized. The H2338 analysis also showed that the resulting structural hypothesis could be connected to a possible functional mechanism, extending the output beyond a list of predicted residues.

The epitope hypotheses were used directly in complex modeling. Full-patch and Top5 restraints generated alternative structure pools, and the router compared those models with the unsteered baseline using the retrieved evidence, predictor confidence, and interface geometry. In H2321, this process recovered a pose that satisfied an escape-derived contact constraint missed by the highest ipSAE baseline. In H2346, it identified an incorrect Fab pairing in the baseline and selected a steered model consistent with both the anti-κ evidence and the expected chain arrangement. Across the target set, the mean DockQ of the final APEs predictions reached 0.375 when the best of three independent selection calls was considered, compared with 0.306 for the unsteered baseline. Among the six CASP17 targets with matched human group (CSSB-human) submissions, the corresponding mean DockQ for APEs was 0.418, compared with 0.261 for the Model 1 submissions from the human CASP group and 0.363 when the best of five human submissions was considered for each target (Fig. S2). Together, these results show that APEs can carry antibody-specific evidence from epitope inference through guided structure generation and final model selection, producing predictions that are competitive with expert CASP submissions.

The results also define the information supplied by an epitope hypothesis. It constrains the antigen surface involved in binding but does not fully determine the antibody orientation. H1225 had a correctly localized epitope, yet neither the steered models nor the unsteered baseline recovered an accurate complex. H2346 exposed a related selection problem: both steered candidates agreed with the epitope evidence, and the router selected the clash-free Top5 model, although the full-patch model had a higher retrospective DockQ. Biological consistency and interface checks were effective for rejecting a clearly incorrect baseline, but a more plausible structural score is still needed to rank plausible alternatives.

This study included only 14 CASP targets, which limits conclusions about performance across broader antibody and antigen classes. We also did not systematically ablate retrieval, agent roles, language-model choice, or reasoning settings, so the contributions of individual components remain unresolved. The date cutoff applied only to external sources retrieved during each run; it could not remove relevant knowledge that the language model may have encountered during pretraining. Additionally, we were not able to obtain epitope predictions for four targets (H2340, H2354, H2355, and H2419) due to the biological safety filter of the models repeatedly interrupting the prediction runs. Evaluation on larger and more diverse benchmarks, including FoldBench^5^ and PXMeter^6^, will be needed to test whether the results generalize beyond the current target set. Within this scope, the study shows that the complete workflow can retrieve antibody-specific evidence, convert it into structural restraints, and use those restraints for model generation and selection.

The agentic design also allows the prediction loop to be extended beyond selection from a fixed pool. When the available structures do not satisfy the evidence, the router could retain competing models, request additional sampling, or revise the steering residues. Evaluating such an adaptive workflow will require prespecified stopping and selection rules, but the present results provide the evidence-to-structure pipeline needed to support it.

## Methods

### Inputs provided to the APEs

We provided the agent with separate antigen and antibody structures, sequence-based chain role assignments, and precomputed database search results through read-only files in its working directory. The antibody-antigen complex predicted by Protenix-v1^3^ was split into two PDB files, one containing the antigen and the other containing the antibody. Each structure was centered independently so that the agent could not use their predicted relative placement. To prevent the agent from using Protenix-derived confidence information, all pLDDT values were set to 0.0. Chain roles were assigned by detecting immunoglobulin variable domains with ANARCI^25^. For each chain sequence, we performed a UniProt^26^ BLAST^27^ search and retained the top 10 hits, together with the corresponding homologous sequences, UniProt annotations, and associated publications. InterPro domain annotations^28^, ANARCI/IMGT^29^ region labels, and up to 20 hits from a BLAST search of each query sequence against the Pataa patent database were also precomputed.

### Epitope hypothesis launchers

We ran three independent epitope hypothesis launcher agents (Codex gpt-5.6-sol/high) to reduce dependence on a single stochastic reasoning and web-retrieval trajectory. Each launcher received the molecular inputs and precomputed search results described above. The prompt first asked the agent to identify the antigen and describe its biological function. The agent was then asked to examine the antigen surface together with the antibody CDR sequence and shape and propose up to four candidate epitope patches. For every antigen residue included in a patch, the launcher assigned a score from 0 to 1 for predicted CDR contact and classified the contact as heavy-chain, light-chain, or combined. The required output included the antigen identification, antibody features, candidate patches and their rationales, supporting and conflicting evidence, regions that had been considered and ruled out, predicted contact classes, and residue-level contact scores. The three launchers could return up to 12 hypotheses in total.

### Epitope gathering and ranking

The highest-confidence proposal from one launcher was not always the strongest hypothesis across the three runs. We therefore passed all launcher outputs to a gathering stage consisting of a geometric organizer, a Refiner agent, and a Judge agent. The Refiner used Codex gpt-5.6-sol/xhigh, and the Judge used Codex gpt-5.6-sol/high.

The geometric organizer grouped patches that referred to the same antigen-surface region. The spatial distance between patches P and Q was defined as following:

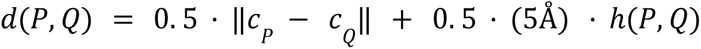

where *c_P_* is the exposure weighted centroid of patch P, and ℎ(*P*, *Q*) is the minimum hop count between P and Q on a residue contact graph G. Two residues were connected in G when any pair of their atoms was within 5 Å. Patches were clustered by single linkage, with the distance between two groups defined as the smallest *d*(*P*, *Q*) among all patch pairs across the groups. Beginning with the closest pair, the organizer merged groups when their distance was below 12 Å and the spatial extent of the combined patch remained within the estimated CDR reach. CDR reach was defined as the 95th percentile of distances from its CDR atoms to their CDR centroid in the input antibody structure.

The Refiner received the organized patches together with the full reasoning logs from all three launchers. Its prompt asked it to review the evidence for each region, develop the strongest biological case supported by the available information, and revise the candidate residue sets where needed. The Judge then received the refined candidates and their associated evidence. It was asked to compare the candidates and assign a relative patch-confidence score according to the evidence that each region was the true epitope. Patch-confidence scores were normalized to sum to 1 within each target and were used to rank alternative hypotheses for that target.

### Evaluation settings and date-cutoff control

To approximate the information available during each CASP experiment, web retrieval was restricted to sources released before the target’s CASP entry date. For each search query, the main agent delegated retrieval to three Codex gpt-5.6-terra/high subagents. Each search subagent returned the retrieved content, its source URL, and a handle used to verify the source release date. Dedicated fetch tools were provided for the RCSB Protein Data Bank^30,31^ and the European Patent Office. Before a search result was passed to the main agent, a code-based gate compared its verified release date with the target-specific cutoff and removed results released after that date.

Epitope predictions were evaluated against reference epitopes derived from the experimental antibody–antigen structures. An antigen residue was included in the reference epitope if any of its heavy atoms was within 5 Å of any antibody heavy atom. The same contact definition was used to derive epitopes from predicted complexes. Residue-level precision was used to measure the fraction of predicted epitope residues that belonged to the reference epitope.

### Epitope-guided sampling and router selection

For each target, the highest ipSAE model among 200 unsteered Protenix-v1 predictions was retained as the baseline. When the rank-1 patch had a relative patch-confidence score of at least 0.4, that patch was used for steering. When its score was below 0.4, each retained candidate was sampled separately. For every candidate, our in-house steering method generated ten seeds with five structures per seed under two restraint settings. The full-patch setting used all residues in the predicted patch, whereas the Top5 setting used the five antigen residues with the highest residue-level contact scores.

We excluded predictor confidence scores for steered structures and selected finalists using contact convergence computed from their final coordinates. For each structure, antibody–antigen contacts were defined as residue pairs with at least one pair of heavy atoms within 5 Å. Contact convergence was the mean Jaccard similarity between the contact set of that structure and the contact sets of the 45 structures generated from the other nine seeds for the same candidate and steering setting. For each target, we retained the model with the highest contact convergence from the full-patch samples and the model with the highest contact convergence from the Top5 samples. The router therefore compared three structures: the unsteered baseline, the full-patch finalist, and the Top5 finalist.

The router agent (Codex gpt-5.6-sol/high) was run independently three times for each target. Each call received the same structure coordinates, interface geometry, steering information, and recorded epitope evidence. Predictor confidence was provided only for the unsteered baseline. The prompt allowed three decisions: **KEEP**, which retained the baseline as the preferred model; **SWITCH**, which selected one of the two steered finalists; and **NEITHER**, which retained the baseline as a fallback without endorsing any of the three structures. The router received no worked examples, previous router decisions, native structure, or web access. All decisions were frozen before whole-assembly DockQ was calculated. We report the decision from each call and, as a retrospective summary, the highest selected DockQ across the three calls for each target.

## Acknowledgements

This work was supported by the National Research Foundation of Korea [RS-2024-00397865] and the Institute of Information & communications Technology Planning & Evaluation (IITP) [RS-2023-00220628] funded by the Korea government (Ministry of Science and ICT). This work was also supported by Korea Health Industry Development Institute (KHIDI) [RS-2025-25461991] funded by the Korea government (Ministry of Health and Welfare)

## Notes

### Competing Interest Statement

The authors have declared no competing interest.

