## Supplementary Material for "APEs: Agentic Prediction of Epitopes to Guide Antibody–Antigen Structure Prediction"

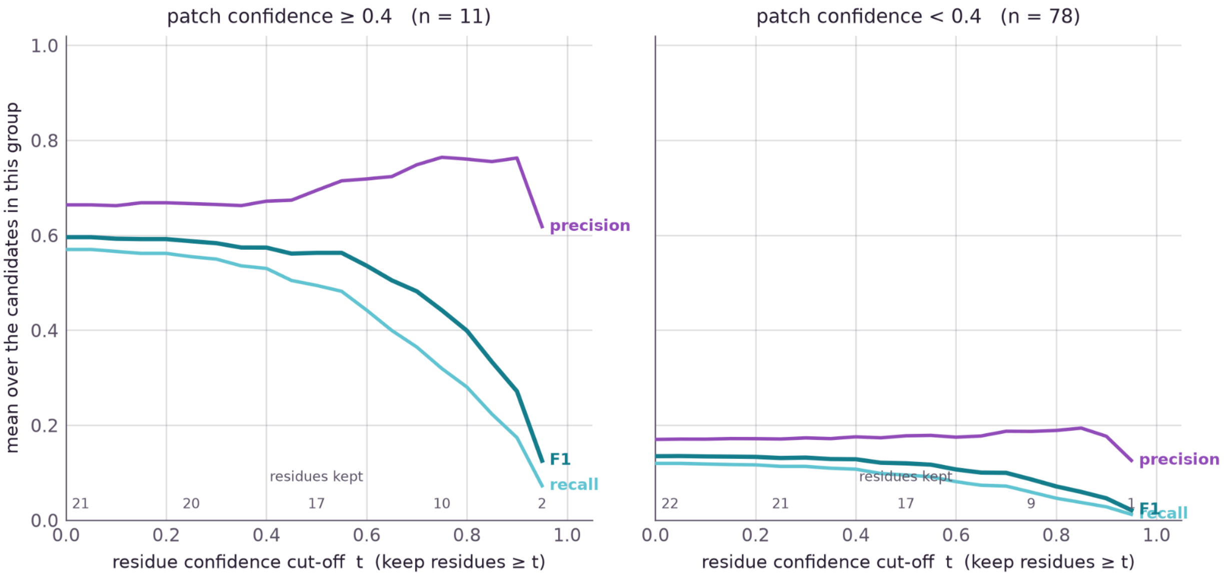


**Figure S1.** Epitope prediction performance across residue-confidence thresholds, stratified by patch confidence

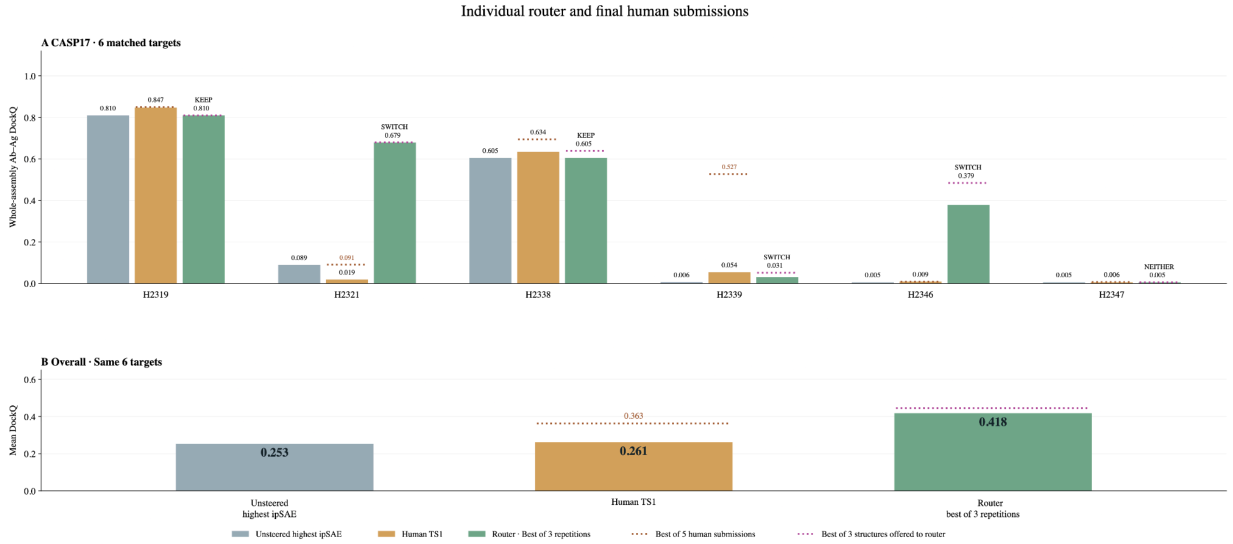


**Figure S2.** DockQ Comparison of highest confidence unsteered Protenix-v1 (gray), Model 1 of a Human Expert CASP predictor (CSSB-Human, yellow), and APEs (green).
